# PREMISE: A database of 20 *Macaca Fascicularis* PET/MRI brain imaging available for research

**DOI:** 10.1101/2023.03.03.530981

**Authors:** Lucie Chalet, Justine Debatisse, Oceane Wateau, Timothe Boutelier, Marlène Wiart, Nicolas Costes, Ines Merida, Jérôme Redouté, Jean-Baptiste Langlois, Sophie Lancelot, Christelle Léon, Tae-Hee Cho, Laura Mechtouff, Omer Faruk Eker, Norbert Nighoghossian, Emmanuelle Canet-Soulas, Guillaume Becker

**Affiliations:** CarMeN Laboratory, Université Claude Bernard Lyon 1, INSERM U1060, INRA U1397, Lyon, France; Olea Medical, La Ciotat, France; Institut des Sciences Cognitives Marc Jeannerod (ISCMJ), UMR 5229 CNRS, Bron Cedex, France; Cynbiose SAS, Marcy-L’Etoile, France; CERMEP, Lyon, France; Hospices Civils de Lyon, Lyon, France; CREATIS, CNRS UMR-5220, INSERM U1206, Université Lyon 1, INSA Lyon Bât. Blaise Pascal, 7 Avenue Jean Capelle, Villeurbanne 69621, France

**Keywords:** Neuroimaging, Non-human primate, data sharing, Magnetic Resonance Imaging, Positron Emission Tomography, brain imaging data structure

## Abstract

Non-human primate (NHP) studies are unique in translational research, especially in neurosciences and neuroimaging approaches are a preferred method for scaling cross-species comparative neurosciences. In this regard, neuroimaging database development and sharing are encouraged to increase the number of subjects available to the community while limiting the number of animals used in research. We present here a simultaneous PET/MR dataset of 20 Macaca Fascicularis structured according to the Brain Imaging Data Structure (BIDS) standards. This database contains multiple MRI sequences (anatomical, diffusion and perfusion imaging notably), as well as PET perfusion and inflammation using respectively [^15^O]H_2_O and [^11^C]PK11195 radiotracers. We describe the pipeline method to assemble baseline data from various cohorts and qualitatively assessed all the data using signal-to-noise and contrast-to-noise ratios as well as the median of intensity. The database is stored and available through the PRIME-DE consortium repository.

## Introduction

In the framework of open science, sharing imaging databases offers specific benefits in terms of analytical tools development and validation [1]. According to the FAIR Data Principles, the development of accessible imaging databases will help increase the reproducibility between studies [2]. In this respect, neuroimaging scientists examine their respective policies and practices [3]. This is also valid in pre-clinical research where in-vivo imaging of non-human primates (NHP) holds great potential in comparative biology and biomedical research [4, 5]. NHP neuroimaging databases enables the scaling of cross-species comparative neurosciences and a better understanding of brain regions’ functions in health and disease. Besides, the establishment and sharing of pre-clinical neuroimaging databases complies with the 3R principles, especially reduction through the use of imaging data-sets, and refinement considering that in-vivo imaging is non invasive. Therfore, pre-clinical neuroimaging data sharing associates open science objectives and 3R principles torwards better reproducibility and transparency in research [6]). However, several challenges must be overcome by the NHP research community,. Historically, single-lab imaging protocols and heavy logistic of research studies resulted in data acquisition inconsistency and discrepancy of obtained results [7]. Ultimately, this may lead to impaired appropriate data comparison between research groups. The NHP research community is currently facing a significant challenge due to the scarcity of animals. The worldwide sanitary crisis caused by SARS-CoV-2 severely impacted the already precarious supply chain of these animals, and the drastic price increase might strongly impact biomedical research [8, 9]. In recent years, the NHP research community has moved forward to tackle challenges ahead, most notably the limited availability of data. The PRIMatE Data Exchange (PRIMatE-DE) initiative addresses this challenge by aggregating independently acquired NHP in vivo imaging datasets[10]. Initially intended for magnetic resonance imaging (MRI) data, the community has worked to standardize data collection, with minimal acquisition specifications, and data architecture allowing data sharing within the framework of open science [10]. The collaborative work allowed us to gradually improve our neuroimaging studies to human standards. In this context, the Brain Imaging Data Structure (BIDS) became the gold standard for organizing and sharing neuroimaging datasets [1]. Nuclear imaging specialists jumped on board and published guidelines to improve the accuracy and sharing of PET data [11, 12, 13]. Therefore, while PET neuroimaging databases exist, their counterparts in NHP are either missing or not easily available. Furthermore, considering the development of hybrid PET/MR scanners and their translational potential in NHP neuroimaging studies, there is a growing need for NHP PET/MR hybrid imaging databases.

We developed a multi-modal database of Macaca fascicularis, acquired on a clinical PET/MR scanner, and constructed with MRI, [^15^O]H_2_O and [^11^C]PK11195 PET images to BIDS standard. The data was formatted using a self-designed python script compensating for missing metadata frequently encountered with pre-clinical and retrospective data. The designed script is available in open-access on GitHub. The entire dataset is available upon request from PRIMatE-DE repository.

## 1 Materials and Methods

### 1.1 Animal cohort description

This dataset, which includes 20 mature male cynomolgus macaques (Macaca fascicularis), was generated using baseline images from primate stroke model described by Debatisse et al. in 2021 [14]. The experimental protocol was approved by the Animal Welfare Body of Cynbiose and the Ethics Committees of VetAgro-Sup and CELYNE CEEA n°42, and was carried-out in accordance with the European Directive 2010/63/UE and ARRIVE guidelines (Animal Research: Reporting in Vivo Experiments).[15, 16]

The subjects underwent combined PET-MRI acquisitions following anaesthesia induced by IM injection of ketamine (4 mg/kg; KetamineVR 1000, Virbac, France) and midazolam (1.3 mg/kg; MidazolamVR 5 mg/mL, Mylan, France). Sevoflurane (1%, variable depending on the animal’s anesthetic depth; SevoFloVR, Abbott Laboratories, France) maintained anaesthesia during acquisition. Animals were intubated and monitored through heart and respiratory rate, ET-CO2, systolic, diastolic and mean arterial pressure, oxygen saturation and body temperature.

### 1.2 PET-MRI acquisitions

PET-MRI sequences were acquired on fully integrated hybrid Biograph mMR PET-MRI 3T Siemens scanner (Siemens Healthcare, Erlangen, Germany). Imaging acquisitions were conducted between 2016 and 2019 with software versions “syngo MR B20P” and “syngo MR E11”.

MRI sequences are described in table 1 with corresponding parameters according to years of acquisition.

**Table 1:**
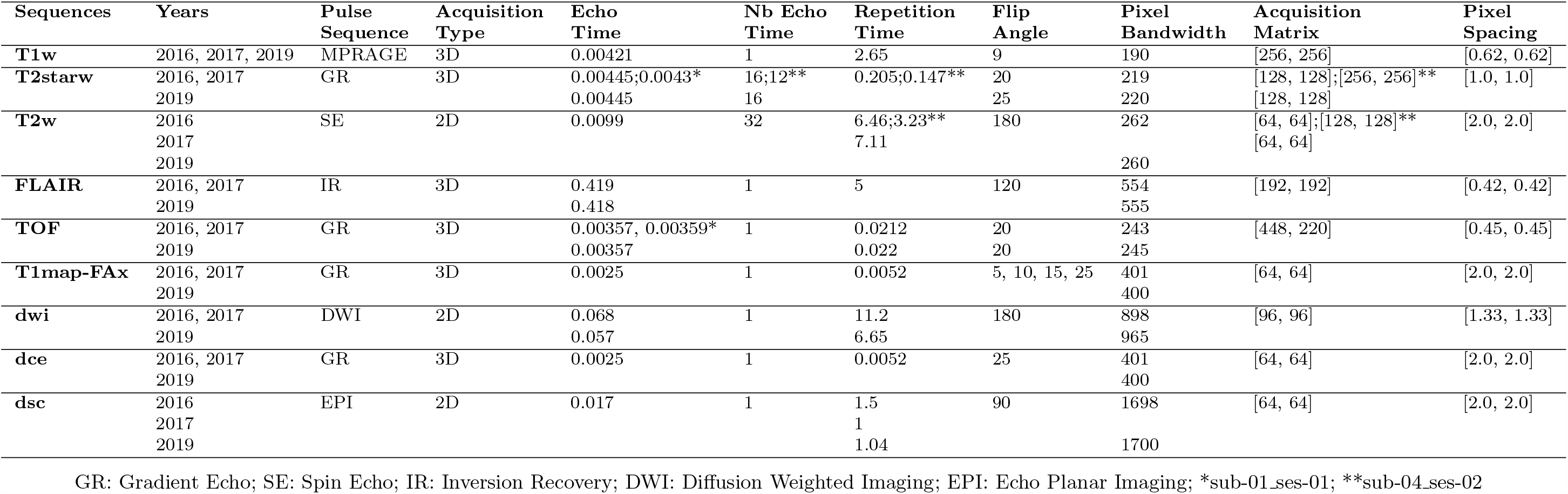
Parameters of MRI sequences

Images from PET radiotracers [^15^O]H_2_O (255 *±* 15*MBq*) and [^11^C]PK11195 (140.1 *±* 21.4*MBq*) were acquired for 6min and 70min respectively, after bolus injections. While molar activity could be measured for [^11^C]PK11195 (48.0 *±* 24.5*GBq/μmol*) providing information on injected mass (3.50 *±* 1.7*nmol*), the half-life of [^15^O]H_2_O did not permit such precise radioactivity measurements. The data were reconstructed on a 256 × 256 × 127 matrix (voxel size: 0.7 × 0.7 × 2.0 mm^3^), 26 cm axial FoV using a point-spread function and OP-OSEM iterative reconstruction method including normalization as well as correction for attenuation, scatter, random counts and dead time. Prior to the PET-MRI session, a CT scan (Siemens Biograph mCT64, Siemens Healthcare, Erlangen, Germany) was obtained for each animal and used for PET attenuation correction. [^11^C]PK11195 dynamic PET data were reconstructed in 28 frames: 6×10s, 6×20s, 6×120s and 8×300s. [^15^O]H_2_O dynamic PET were reconstructed in 26 frames: 8×4s, 4×6s, 6×10s, 8×20s. Lastly, a post-reconstruction 3D gaussian filter of 4mm was applied.

### 1.3 BIDS formatting

The data processing pipeline consisted in formatting the datasets to Brain Imaging Data Structure (BIDS) [1]. Existing formatting tools such as *dcm2bids* [17] have limited support for missing DICOM tags frequently encountered in preclinical and retrospective datasets. Therefore, an automated python script compatible with MRI and PET acquisitions was developed. Raw images are loaded in DICOM format and converted to NIfTI format with associated metadata in json format in accordance with BIDS guidelines. An overview of the python script tasks is provided in figure 1. The script’s ambition is to provide flexibility in data organization and selection when formatting raw data for sharing purposes. Because raw-data folder organization varies between structures, additional parameters are provided enabling the definition of the raw folder structure. For instance, single session studies are supported by the pipeline. It is also possible to select only baseline acquisitions or anatomical images to be formatted to BIDS. This option may be useful in multi-phase studies in which data exploration of key-phases delays data sharing of baseline acquisitions. Additionally to the data conversion, the script provides templates for mandatory BIDS files such as dataset description and participants list.

**Figure 1:**
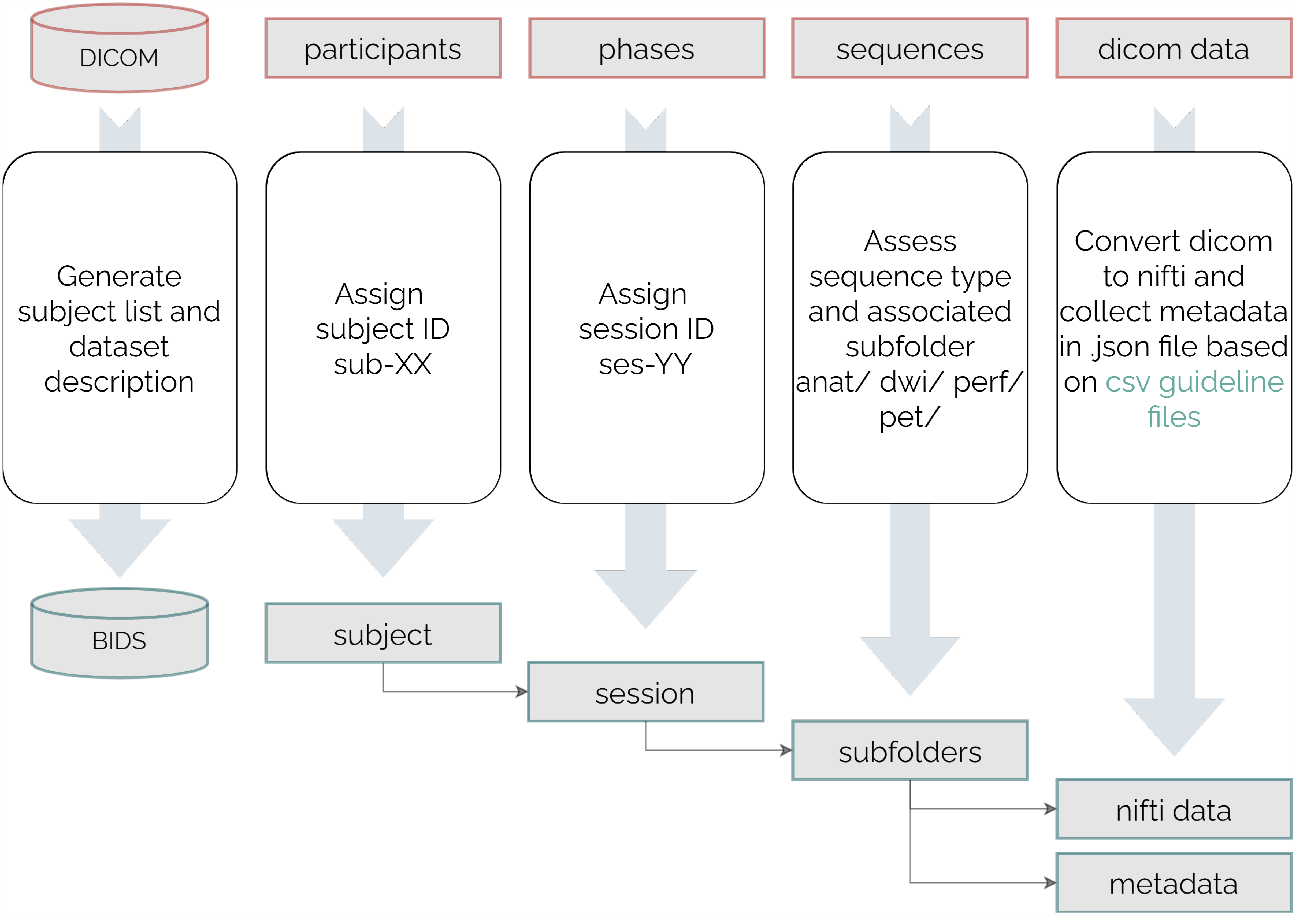
Processing pipeline tasks overview to convert raw DICOM structure to BIDS structure.

#### 1.3.1 Converting pixel data to NIfTI image

DICOM volume loading and conversion to NIfTI format is performed using the *pydicom* and *nibabel* python packages [18, 19]. These packages were integrated in a python class handling multi-dimensional volumes from lists of DICOM files enabling the building of pixel data arrays and managing data orientation to improve NIfTI encoding. Limitations remain for oblique orientations, frequently encountered in large animal models, special care should be provided when handling NIfTI conversions and manual corrections might be necessary. Therefore, the orientation of the pixel data was manually validated using the ITK-snap software [20]. The provided data were acquired on a clinical PET-MRI system without stereotaxic frame, therefore a unique subject position was established close to a patient position. The oblique encoding of MRI sequences led to potential errors in conversions. The closest orientation was Right-Superior-Anterior (RSA) for the majority of subjects. Other subjects were oriented oblique closest to Right-Anterior-Inferior (RAI). In order to provide uniform data, the identified RAI subject’s data were manually oriented to RSA. Additionally, diffusion weighted imaging (DWI) and diffusion tensor imaging (DTI) require bval and bvec files to qualify the sequences. These files are obtained with the *dicom2nifti* python package [21].

#### 1.3.2 Generating the metadata file

To collect the necessary metadata for each acquisition sequence, compensate missing tags and integrate subject-specific radioactivity parameters, three configuration files are necessary.

##### Sequence overview

The sequence overview configuration file provides a list of sequences to integrate in the BIDS formatting. It provides details on tags necessary for each sequence and replacement values if the tag is missing in the DICOM metadata. In this configuration file, the naming format for the sequence in the BIDS database is also defined along with instructions on where to find and store the corresponding data. Figure 2 provides a description of each column required in the sequence overview configuration file along with examples. Each tag is attributed a value depending on the sequence-specific requirements, the tags availability and format in the DICOM metadata and the modality:

**Figure 2:**
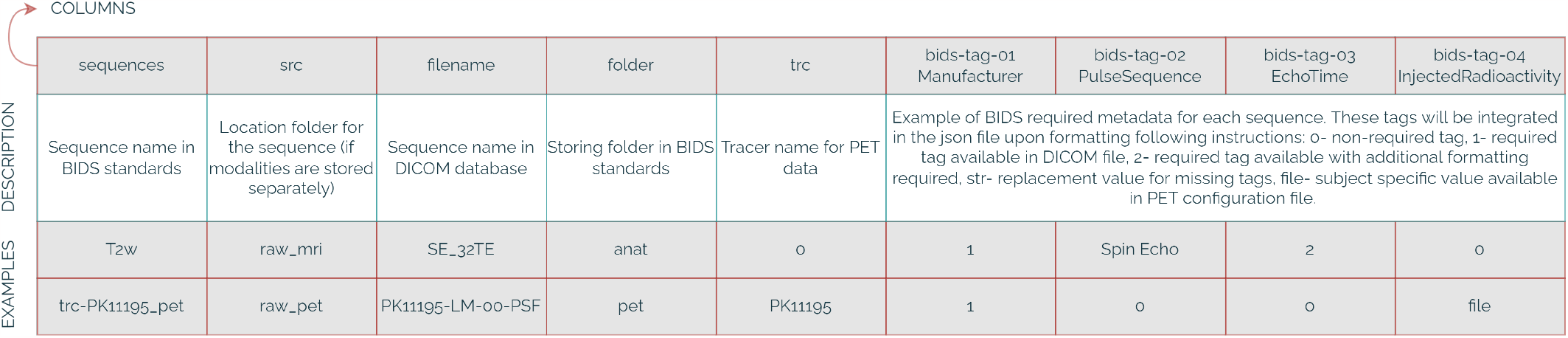
Sequence overview configuration file description and examples.

- 0- Tag not included in the json metadata file of the sequence.
- 1- Tag included in the json metadata file of the sequence as an exact copy of the equivalent DICOM tag.
- 2- Tag included in the json metadata file of the sequence as a formatted copy of the equivalent DICOM tag.
- str- Tag value manually set for the json metadata file due to a missing value in the DICOM file.
- “file”- Variable tag throughout subjects which value must be extracted from “PET doses” file.

##### PET doses

The PET-specific configuration file enables the definition of each subject’s injected radioactivity parameters. The file sets values for BIDS-guideline required PET tags *SpecificRadioactivity* (= Molar Activity, current term recommended by the guidelines [22]), *InjectedRadioactivity* and *InjectedMass*. This file is only required for PET data formatting. Configuration file description and examples are provided in Figure 3.

**Figure 3:**
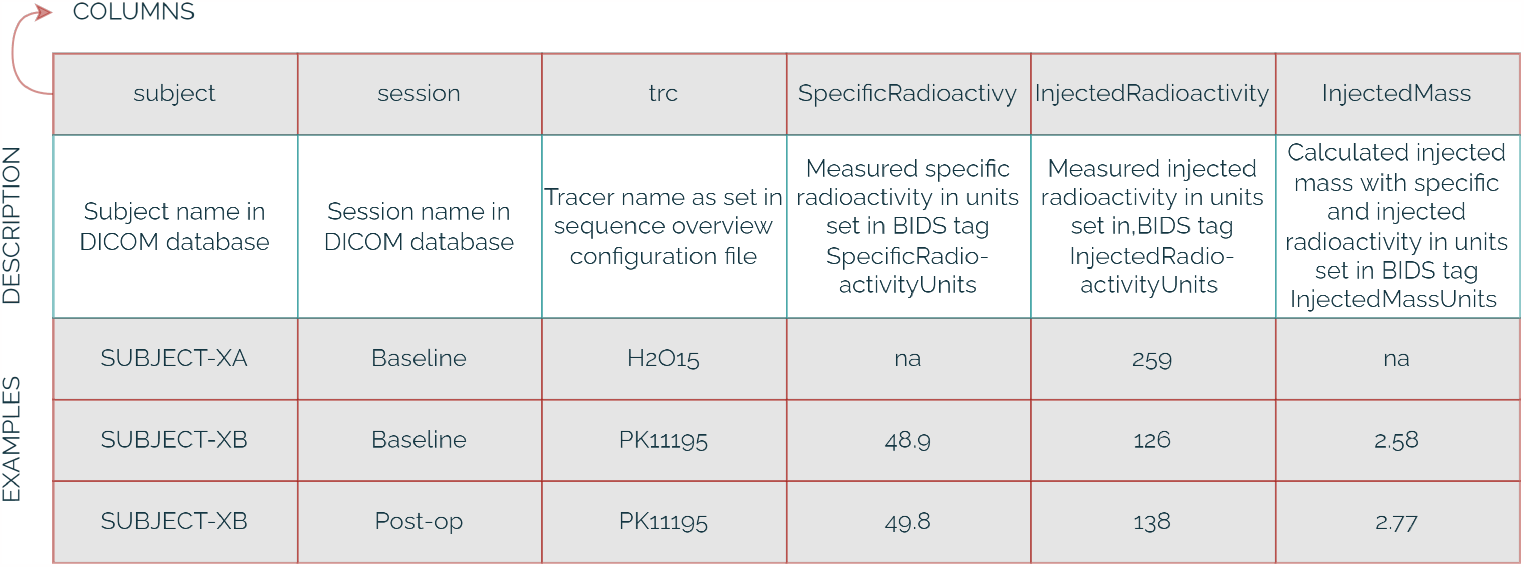
PET doses configuration file description and examples.

##### DICOM to BIDS tag converter

This configuration file provides the DICOM tag name for each required BIDS tag compensating for variable naming convention between the two formats. If no DICOM tag equivalent exists for a given BIDS requirement, the sequence overview or PET-doses replacement option is triggered. Description and examples of the configuration file are given in Figure 4.

**Figure 4:**
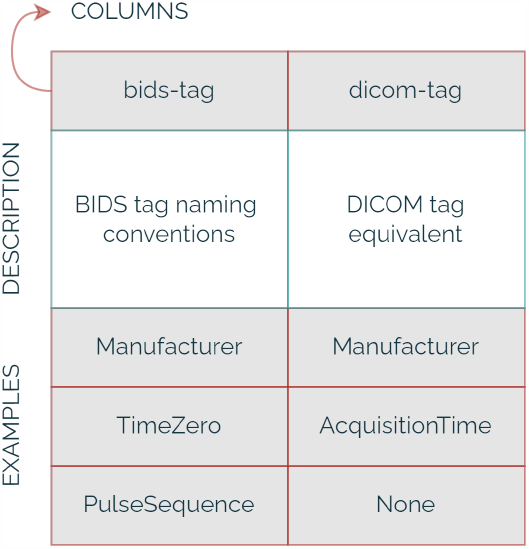
DICOM to BIDS tags conversion configuration file description and examples.

### 1.4 Data quality assessment

To provide an indication of the quality of the database formatted to BIDS standards, we analyzed the distribution of three relevant metrics: signal-to-noise ratio (SNR) and contrast-to-noise ratio (CNR) for MRI acquisitions, and median intensity for PET and MRI acquisitions. The metrics were measured on voxels with tissue signal (volume of interest, VOI) following Otsu thresholding on the volume with highest intensity [23]. Voxels below the given threshold were considered background for noise measurement. The metrics were calculated using the following formulas:

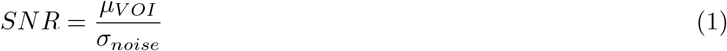

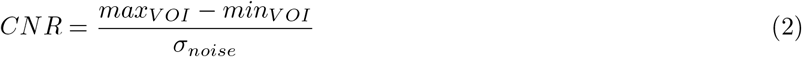

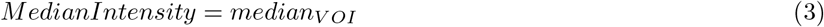

Categorical violin plots were plotted for each metric. These plots show the distribution of the metric values across subjects. Strip-plots were superimposed to the violin plots to provide additional information on time-distributions. In these plots, each category corresponds to an MRI sequence or PET tracer, and the points represent the metric values for each acquisition within that sequence. For dynamic MRI sequences and PET acquisitions, we averaged the metrics across time points or frames and plotted the resulting values. For multi echo sequences, we averaged the median of intensity across echo times and plotted the resulting values. In terms of noise evaluation, only the first echo was plotted as signal is maximal at this echo time. A sample of the data was manually inspected to ensure that the results were accurate and to identify any issues that might have been missed. Based on these evaluations, we were able to provide an indication of the overall quality of the database for potential future users of the dataset, including the detection of potential outliers and the assessment of the homogeneity and consistency of the data.

## 2 Results

After converting the data to BIDS standards, the sequences’ availability was assessed using an automated counter. The data availability is provided with Figure 5. This analysis highlighted the presence of a test-retest for two subjects in the dataset and a few missing acquisitions. The lack of data is explained by issues with radio-synthesis, contrasting agent or movement during acquisition for PET, contrast imaging and MRI sequences respectively. This figure also indicates the study year for each subject, ranging from 2016 to 2019. This parameter is relevant to explain variability in data quality due to software updates on the PET-MRI system.

**Figure 5:**
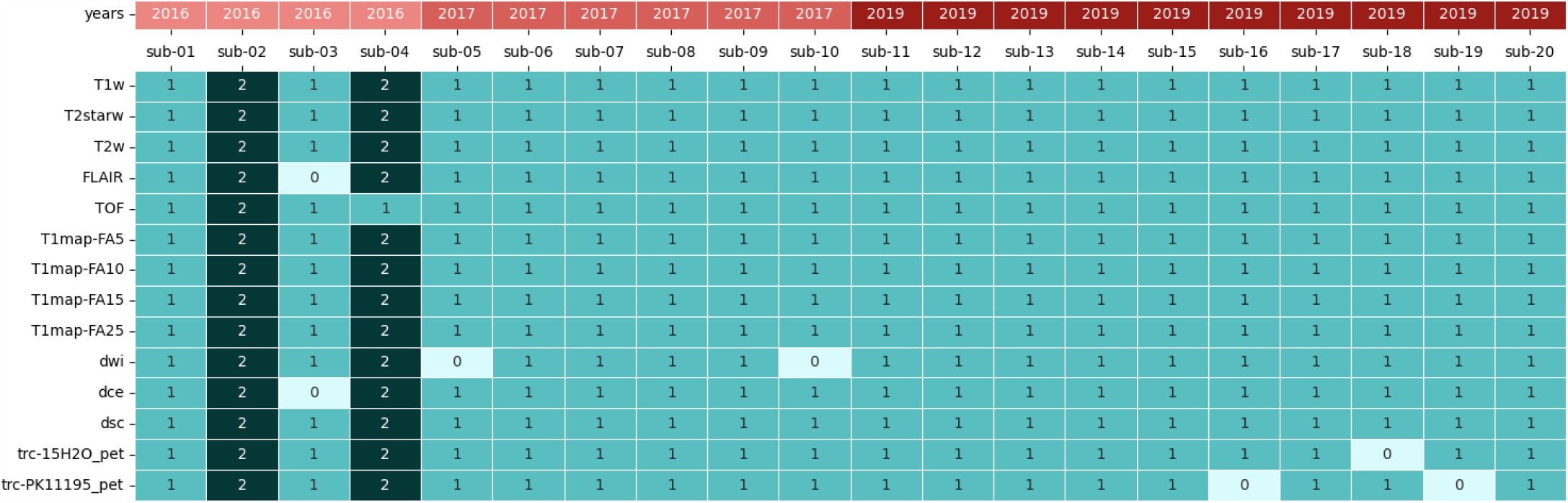
Data availability in shared BIDS database with study years for each subject.

As previously described, the quality of the acquisitions in the shared BIDS database is expressed with SNR, CNR and median of intensity. The quality assessment highlighted the consistency of the acquisition quality (averaged and over time) as displayed in Figure 6.

**Figure 6:**
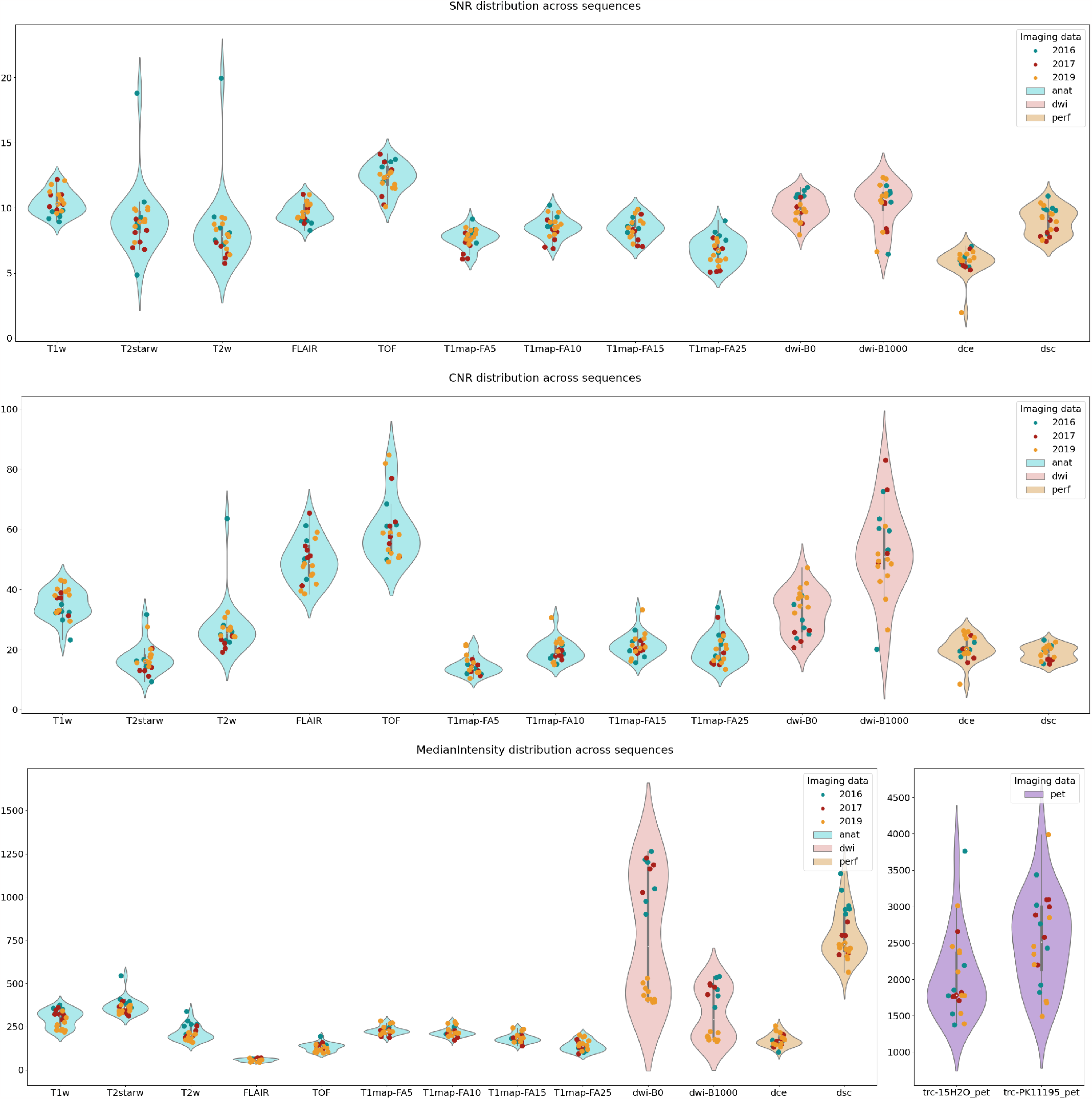
PET and MRI acquisitions quality compared between subjects across sequences according to SNR (top), CNR (middle) and median of intensity (bottom).

While acquisition quality was highly reproducible, an outlier in T_2_-weighted and T_2_^*^-weighted sequences was identified. This outlier is due to a variation in acquisition parameters as shown in table 1. The DCE-perfusion acquisitions also uncovered an outlier due to the movement of the identified subject 15 during acquisition.

Two groups can be discriminated in the DWI metrics evaluation. This distinction is due to a software update in 2019 after which the sequence’s parameters were strongly modified for this acquisition. As a result of the software update, the median of intensity is significantly reduced. A variation in signal-to-noise and contrast-to-noise ratios can also be observed, however with milder difference between software versions.

A variability of the PET signal intensity can be observed.

## 3 Discussion

In line with human neuroimaging advances, the NHP scientists community is evolving toward more reproducibility and transparency in biomedical sciences, notably through the means of data sharing. To the best of our knowledge, this NHP imaging database is the first open-source collection including simultaneous dynamic PET-acquisition and MRI sequences. We provide a wide range of NHP MRI acquisitions for download, including structural, diffusion and perfusion imaging.

BIDS standards were originally designed to guide best practices for storage and sharing of fMRI data sets [1]. PET imaging modalities came afterwards as extensions to the original BIDS specifications [12]. In this context, Drs Gitte Knudsen and Robert Innis initiated a collaborative working program to address specific PET imaging challenges through the definition of standards for organizing and sharing [11]. We followed the current recommendations regarding the description of data acquisition and reconstruction methods as well as the molecular description of the radio-tracers. The purpose of the present brain NHP database is to provide the molecular imaging community a full data set with detailed quality description. We included quality control measures of PET images using the median of intensity metric. Although the methods we propose here is derived from MRI applications, we believe that they can be used to quantify the variability of the different PET data acquisition. This variability is due to the injected radioactivity, subject’s weight and physiological constants’ variations. While corrections are applied to normalize weight and dose variations using standardized uptake value (SUV) quantification, variability of physiological constants can be compensated using reference region ratios (SUVr). We provided uncorrected data enabling future users to apply their own normalization and modeling methods.

We are aware of the quality limitations of the MRI data in comparison to the standards in NHP imaging [24]. These drawbacks are largely due to the specifics of our research protocol dedicated to translational stroke research. Therefore, baseline acquisition (before stroke induction) were acquired in the same conditions as further occlusion-reperfusion acquisitions [14, 25] which precludes the use of a stereotaxic frame for a repeatable animal position in the scanner. Moreover, due to the specificity of our model, acquisitions had to be shortened to fit experimental conditions in stroke phases. While we observed variability in our data quality which might alter the automation of pipelines; the diversity of our database can represent a source of interest for a wide range of applications from noise reduction to anatomical studies.

When formatting a retrospective and/or pre-clinical database to BIDS standards, we frequently encounter missing data and missing DICOM tags. These inconsistencies are difficult to identify and compensate. Therefore, we found a need for a self-designed tool to format data following BIDS guidelines. Additionally, this tool enables the selection of specific setting/experimental phase to include in the converted database. We found this functionality could encourage scientists to share parts of their data while holding the remaining settings until results are published. We also support a variety of raw data folder organization as we know it varies between structures and institutes. A common issue faced in the development of data formatting and sharing is the time and resources these initiatives require. This highlights the need for data scientists in research teams dedicated to these tasks. We are hoping that the provided tool will facilitate such initiatives that are urgently needed to address societal challenges raised by the use of NHP in biomedical research.

To conclude, we generated an original and diverse NHP hybrid PET/MR database available for the community through PRIME-DE platform and hope that the present note, describing the quality of the published data and metadata will encourage the neuroimaging community to use it.

## 4 Acknowledgments

This work was supported by ANR CYCLOPS and CMRO2 (ANR-15-CE17-0020 and ANR-21-CE17-0028), the RHU MARVELOUS (ANR-16-RHUS-0009) of Lyon University, under the Investissements d’Avenir program of the French National Research Agency (ANR). The Ph.D. salary of LC (Cifre, OLEA Medical) is co-funded by the French Ministry of Higher Education and Research (ANRT).

